# metaPathwayMap: A tool to predict metabolic pathway neighborhoods from structural classes of untargeted metabolomics peaks

**DOI:** 10.1101/2022.03.15.484337

**Authors:** Gaurav Moghe, Susan Strickler

## Abstract

**Summary:** Thousands of peaks detected via untargeted tandem liquid chromatography mass spectrometry (LC-MS/MS) of natural extracts typically go unannotated, limiting our understanding of the metabolic pathways perturbed under different conditions. Current tools for predicting metabolic pathways from untargeted metabolomics data either require prior compound identification or are more focused on specific model species. metaPathwayMap makes use of recent advances in computational metabolomics to map peaks detected in untargeted LC-MS/MS experiments to MetaCyc pathway representations using their structural class predictions. This approach enables better insights into metabolomes of model and non-model species.

**Availability and Implementation:** Required Python scripts can be downloaded from the moghelab/metaPathwayMap GitHub repository and implemented on a Unix machine. This tool is also available for use through the SolCyc website (https://metapathwaymap.solgenomics.net) and via DockerHub (srs57/metapathwaymap).

**Supplementary Information:** Additional information is provided in Supplementary Methods, Supplementary Files 1-3 and on GitHub (https://github.com/moghelab/metaPathwayMap)

## Introduction

Organisms respond to environmental stress by modulating flux through their complex metabolic pathways. Untargeted metabolomics studies using tandem liquid chromatography mass spectrometry (LC-MS/MS) aim to capture these perturbations, however, of the thousands of peaks detected in natural extracts, <5% can be reliably identified by matches to spectral databases (da Silva *et al*., 2015). Thus, the significance of global metabolome shifts under stress or in mutants remains understudied compared to transcriptomics, where database searches can reliably identify properties such as protein domains, orthologous groups, and Gene Ontology. To circumvent this lack of identifiable peaks, metabolic profiling studies typically target specific compound classes, however, this experimental design ignores global changes occurring in the metabolome. Tools mapping metabolomic data to pathways, such as mummichog (Li *et al*., 2013), MetScape (Basu *et al*., 2017), MetaboAnalyst (Pang *et al*., 2021), metaMapp (Barupal *et al*., 2012) are restricted to a handful of model organisms and mostly require prior compound identification, limiting the study of metabolome perturbations in non-model species. This also leaves >95% of the metabolome unstudied. Thus, new strategies to better predict perturbed metabolic pathways from untargeted metabolomics data are needed.

A recent, deep learning based tool named CANOPUS (Dührkop *et al*., 2020) circumvents the problem of compound identification. CANOPUS, part of the SIRIUS package (Dührkop *et al*., 2019), takes MS/MS data as input, and uses pre-trained deep learning models to predict structural categories (e.g. Superclass, Class, Subclass, Level5, …, Most specific class) representing the likely dominant and other minor structural motifs. These categories are based on the well-established ChemOnt ontology (Feldman *et al*., 2005), which has also been applied to entries in the Chemical Entities of Biological Interest (ChEBI) database (Degtyarenko *et al*., 2008) using the tool ClassyFire (Feunang *et al*., 2016). However, many most-specific annotations of CANOPUS (e.g. cyclitols and derivatives [for shikimate], 5’-deoxy-5’-thionucleosides [for *S*-adenosylmethionine], naphthoquinones [for plumbagin]) are not intuitive/specific enough for downstream pathway or compound identification.

In this study, using the ChemOnt classification of ChEBI entries as a common thread **(Supp. Methods Fig. 1A)**, we developed a novel tool metaPathwayMap (mPm) that matches CANOPUS structural assignments of LC-MS/MS peaks to compounds in the MetaCyc-format pathway database representation (here, PlantCyc) (Hawkins *et al*., 2021). The outcomes are (i) a novel clustering of compositionally similar pathways, which we refer to as “pathway neighborhood”, and (ii) prediction of this structural and pathway “neighborhood” of the metabolomics peak, which is useful in pathway/enzyme discovery. mPm predictions can be used, for example, to generate hypotheses about the importance of differentially abundant metabolites, or to identify relevant pathway enzymes in conjunction with orthogonal RNA-seq data.

## Materials and Methods

mPm uses two inputs: (1) MetaCyc (here, PlantCyc)-format based pathway models and (2) CANOPUS predictions of structural classes of LC-MS/MS peaks. MetaCyc flat files provide compound names, pathways, reaction layouts, and ChEBI IDs for most compounds. Only compounds described in Reaction-Layout descriptor, with ChEBI IDs are considered. Since each compound can belong to >1 pathways, we first constructed a pathway similarity network **(Supp. File 1)** based on the similarity of the ChemOnt categories of the pathway’s compounds. Distance (1-Jaccard Coefficient) was calculated between pathway pairs. The 1^st^ percentile of the distribution of distances from bootstrap comparisons was used as threshold to make clusters. Users can generate pathway representations of other MetaCyc datasets using provided scripts.

mPm further computes Jaccard Coefficients between each CANOPUS prediction and each MetaCyc compound, and filters the predictions using a user-defined threshold. If Compound A is part of Pathways X,Y,Z, all three pathways will be listed as of interest, and the network component to which each pathway belongs is listed to identify compositionally similar pathways. Tab-delimited as well as Cytoscape compatible network files are provided as output.

In addition to the downloadable Python-based code, a GUI-based web application implementing the mPm pipeline was developed using Django v4.0.2 (Django Software Foundation, 2019) and PostgreSQL 14.1. The web application can be accessed at SolCyc (Foerster *et al*., 2018). Version of the web app is also available on DockerHub.

## Results

We tested the accuracy of mPm in predicting the right compound and pathway neighborhoods using **(Set 1)** 17 high-confidence anthocyanins from sweet potato (Bennett *et al*., 2021) and **(Set 2)** 21 mixed compounds from the MassBank database (Horai *et al*., 2010) and additional resin glycosides and acylsugars (Landis *et al*., 2021; Kruse *et al*., 2022) **(Supp. File 2A**,**B)**. CANOPUS annotated 4/17 Set 1 anthocyanins incorrectly as Flavonoid 3-O-p-coumaroyl glycosides (Most Specific Class), while others were annotated as Anthocyanidin 3-O-glycosides. Other classes were consistent for all 17 compounds. Nonetheless, mPm assigned all compounds correctly to different anthocyanin biosynthesis/modification pathways **(Supp. File 3A)**.

Of the 21 Set 2 compounds, 20 received CANOPUS annotations **(Supp. File 3B-D)**, of which the main classes (representing the classes of the largest structural moiety) were correct for only 10 (50.0%). If the alternate classes are considered, then correct class predictions were obtained for 15 (75.0%) **(Supp. File 3B-D)**. Of these 15, PlantCyc pathways of 12 (80.0%) were correctly predicted by mPm **(Supp. File 3B-D)**. Two additional compounds – coniferaldehyde and glutamate – did not map accurately but were assigned to pathways that were first neighbors of the correct pathways in the neighborhood network. Considering these, 14 (93.3%) compounds were correctly mapped to pathway neighborhoods. Pathways of *myo*-inositol were incorrectly predicted as mannitol/crocetin/trehalose biosynthesis. Three compounds – Malvidin-3-O-galactoside, Tricolorin A, acylsugar hab 737 – were not in PlantCyc pathway representations. However, both CANOPUS and mPm identified the correct structural neighborhoods for the first two compounds. Both tools failed for the acylsugar example.

No comparable tool exists for direct comparison of performance. As a proxy, we surveyed the CSI-FingerID (Dührkop *et al*., 2015) annotations in the SIRIUS tool to determine their utility in pathway identification using either “all Bio databases” or specific Natural Product databases **(Supp. Files 3E-G)**. Of the 17 and 14 identified Set 2 compounds, only 7 (41.1%) and 8 (57.1%), respectively, were intuitive for identifying pathways. Thus, by using predictions of both main and alternate structural classes, mPm may be able to build upon the predictions of SIRIUS, CSI-FingerID and CANOPUS to identify structurally similar compounds/pathway neighborhoods. We note that the identifications are dependent on the input pathway representations and high-quality MS/MS data enabling CANOPUS predictions. These predictions, for example, can be further overlaid with RNA-seq data to identify specific enzymes and pathway steps perturbed in control-vs-test comparisons, generating improved functional hypotheses for experimental validation.

## Usage

mPm can be run on Unix using Python scripts downloadable from our GitHub repository (moghelab/metaPathwayMap). This tool is also embedded on the SolCyc website. Only the CANOPUS output, obtainable from the SIRIUS4 package, is needed as variable input for mPm.

## Acknowledgements

We thank Elizabeth Mahood for providing the anthocyanin data, Dr. Lukas Mueller for providing guidance on SolCyc integration, and Dr. A. Daniel Jones for helpful feedback.

