## Supplementary Methods for "metaPathwayMap: A tool to predict metabolic pathway neighborhoods from structural classes of untargeted metabolomics peaks"

#### A. Selection of compounds in Sets 1 and 2

We selected entries in the MassBank of North America database (Horai *et al.*, 2010) representing a diverse set of compound classes including alkaloids, sugars, amino acids, glucosinolates, glycolipids, aminoacyls, phenylpropanoids, indole compounds, sulfur containing compounds and anthocyanins. We manually selected entries with either a CLEAN tag on MoNA or where multiple peaks were observed without obvious noise. No consideration for qTOF or Orbitrap was given. Additional compounds from our own in-house studies on anthocyanins, resin glycosides and acylsugars were also added **(Supp. File 3A,B)**. These had been obtained from Orbitrap (anthocyanins, resin glycosides) and qTOF (acylsugars)(Kruse *et al.*, 2022; Landis *et al.*, 2021). No filtering of any MS/MS peak was performed.

#### B. Running SIRIUS4 and CANOPUS

SIRIUS4 (Dührkop *et al.*, 2019) was run from the Windows GUI. For SIRIUS fragmentation tree calculation, CHNOPS was specified with atoms from 0 to inf. For CSI-FingerID (Dührkop *et al.*, 2015), either “Bio database”, or a set of natural product databases, or only PlantCyc were specified **(Supp. File 3)**. Result summaries were exported without any additional parameter changes, and the tab delimited adducts file for CANOPUS predictions was analyzed. Only class predictions with posterior probability > 0.5 are exported by default.

#### C. metaPathwayMap runs

metaPathwayMap was run on a x86_64 GNU/Linux machine without specifying any memory or extra processors. Detailed instructions for running part 1 (get_similar_pathways_wrapper.py) and part 2 (metaPathwayMap.py) are provided on the moghelab GitHub page. Overall rationale of metaPathwayMap is shown in **Supp. Methods Fig. 1**. The scripts produce several intermediate files that can be used for internal analyses by the user.

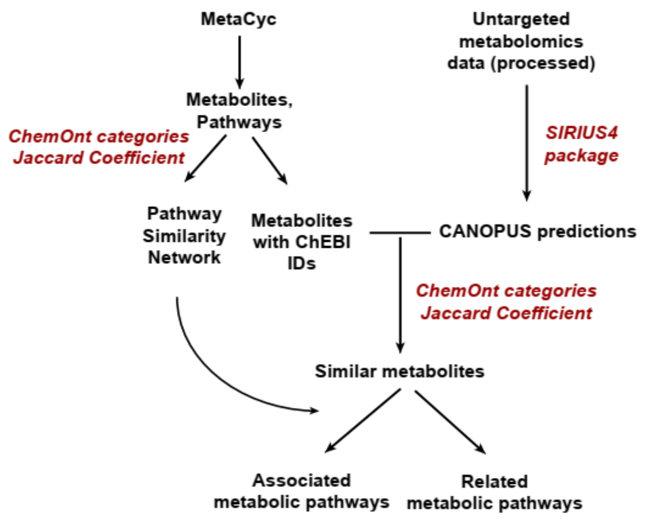


**Supp. Methods Fig. 1:** Overall rationale of the metaPathwayMap pipeline.

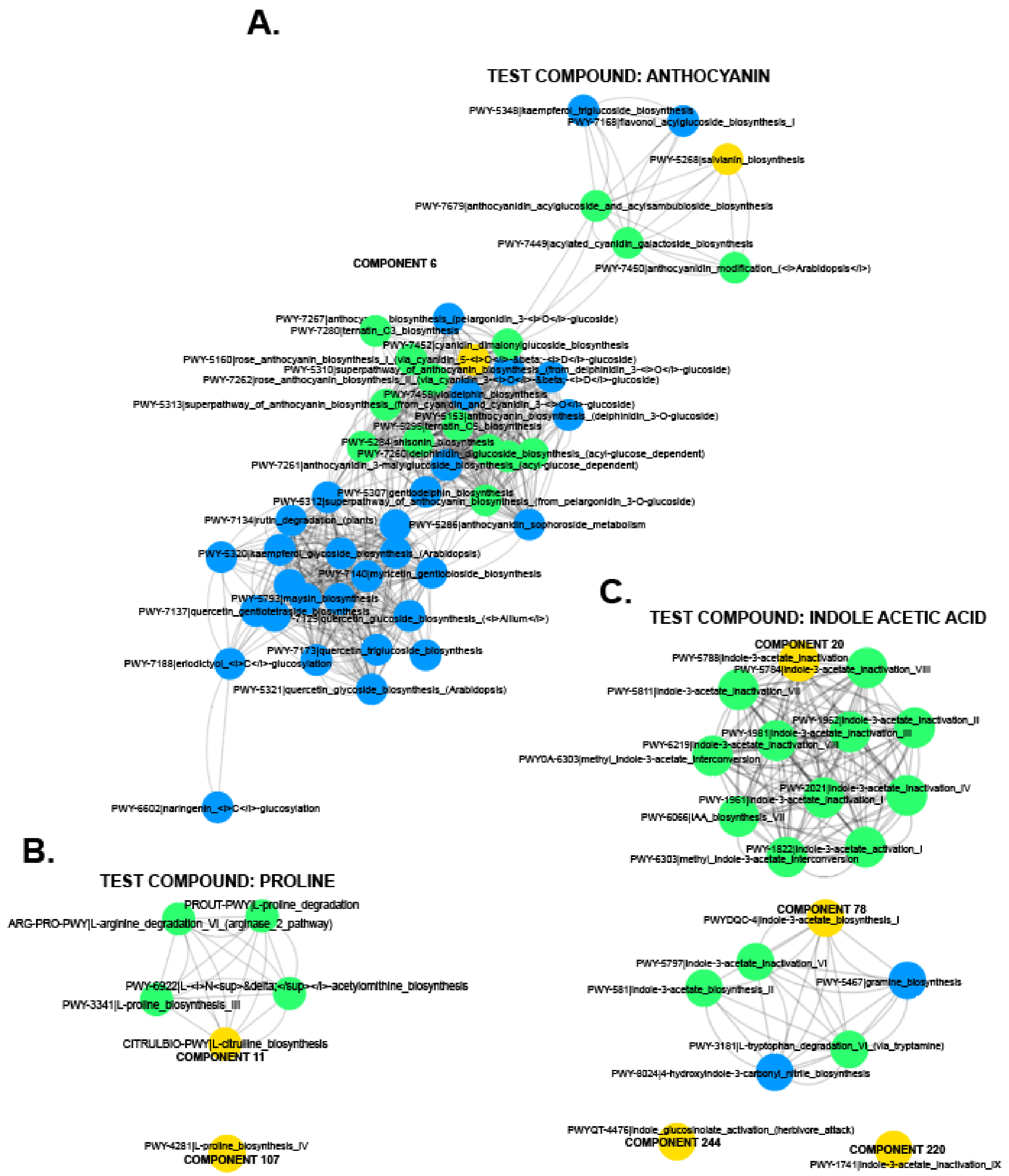


**Supp. Methods Fig. 2:** Three examples of pathway neighborhoods identified from metaPathwayMap. Blue nodes are the basal Cytoscape node colors. Yellow is the first hit shown for a given component by metaPathwayMap. Green nodes are other pathways where the predicted compound is found in, in the main reaction steps.

### D. Interpretation of results and network components

There are four issues to consider when interpreting results:

***(i) Identifying appropriate network components may require different thresholds than the default:*** Empirical testing suggested that the 1st percentile of the random Jaccard scores did not mix unrelated pathways. In some cases, a fixed threshold of 0.20 may be desirable, and both files are provided as outputs by the get_similar_pathways_wrapper.py script.

***(ii) Jaccard similarity scores can be indicative of predictive accuracy:*** metaPathwayMap matches CANOPUS structural class predictions to structural classes of PMN compounds. The top match only shows the most similar compounds thereby linking the pathways associated with it. Empirical observations with the 38 test compounds in Sets 1 and 2 suggest that higher the score, the greater chance that the compound being matched is correct or compositionally very similar. Compound matches with coefficients above 0.7 are generally accurate (with respect to the structural neighborhood), between 0.6-0.7 are more accurate than not, and below 0.6, predictions tend to be noisy. We do not explicitly test these hypotheses in our study or optimize the similarity score (which may, in fact, be class-specific), instead listing all the possibilities for the end user to decide.

***(iii) A compound can be part of many pathways:*** metaPathwayMap identifies up to 5 compounds satisfying the user set Jaccard similarity threshold matching to the provided peak. For each compound, all pathways the compound is associated with in the provided pathway database are listed. metaPathwayMap cannot differentiate between the pathways of a given compound – orthogonal datasets like RNA-seq can be used for that purpose.

***(iv) Matches of a CANOPUS-annotated peak may be incorrect:*** While we show that 80% of our test compounds were matched correctly with their pathway neighborhoods, natural extracts may contain compounds with different structural scaffolds, completely unknown pathways, or pathways not mapped in the PMN databases, which metaPathwayMap will not be able to identify. However, if CANOPUS annotations are correct, metaPathwayMap is highly likely to identify compositionally similar pathways (e.g. Malvidin-3-O-galactoside, Tricolorin A in **Supp. File 3B**).

***(v) metaPathwayMap shows pathway neighborhoods:*** A pathway neighborhood is defined here as a cluster of pathways composed of structurally similar substrates. This is viewed as a network cluster **(Supp. Methods Fig. 2)**. The predictions of metaPathwayMap should thus be considered initial hypotheses, just like the outputs of other predictive software tools, to be later verified through orthogonal means. These may include a focused study of MS/MS patterns to validate compound ID, integrating RNA-seq results or more detailed functional validation studies.

***(vi) metaPathwayMap may be able to distinguish between correct and incorrect CANOPUS predictions:*** metaPathwayMap performs well for compounds for which CANOPUS predicts correct structural classes. However, if CANOPUS predictions are completely wrong, then metaPathwayMap predictions will also be wrong. Nonetheless, we found that in five cases in Set 2 where CANOPUS annotations were completely incorrect, no metaPathwayMap predictions >0.6 were provided for three of those compounds, and for one additional compound, the score was 0.61. Thus, metaPathwayMap may mitigate the effects of incorrect CANOPUS predictions. Clean and good quality MS/MS fragmentation is helpful for both CANOPUS and metaPathwayMap.
